# Complete mitochondrial genome and draft chloroplastic genome of Northern Atlantic *Haslea ostrearia* (Gaillon/Bory) Simonsen, 1974 (Naviculaceae, Bacillariophyceae)

**DOI:** 10.1101/2022.11.09.515767

**Authors:** Aurelie Peticca, Mostefa Fodil, Helene Gateau, Jean-Luc Mouget, Francois Sabot, Benoit Chenais, Nathalie Casse

## Abstract

The first completed, circular mitochondrial genome and the first draft chloroplastic genome of Northern Atlantic blue diatom *Haslea ostrearia* (Naviculaceae, Bacillariophyceae) are described. The mitochondrial genome is composed of 38,696 bases and contains 64 genes, including 31 protein-coding genes (CDS), 2 ribosomal RNA (rRNA) genes and 23 transfer RNA (tRNA) genes. For the chloroplast, the genome is composed of 130,200 bases with 169 genes (131 CDS, 6 rRNA genes, 31 tRNA genes, and 1 transfermessenger RNA (tmRNA) gene). Phylogenetic trees suggest the proximity of all *H. ostrearia* strains yet available and the possibility to use these genomes as future references.

---

*Haslea ostrearia* is a blue microalga from the Naviculaceae family, which lives freely in benthic marine environments or as an epiphyte on brown macroalgae (Simonsen 1974). The blue colour comes from a pigment called marrenine that *H. ostrearia* produces and accumulates at cell apices (Gastineau et al. 2014A). This specific pigment is responsible for the greening of oyster gills in farming ponds in Western France. Furthermore, it has been shown that marrenine displays antibacterial, antiviral and antifungal effects (see Gastineau et al. 2014b). The whole DNA of a *H. ostrearia* originating from the Northern Atlantic Ocean was sequenced, allowing the reconstruction of its mitochondrial and chloroplast genomes. This study clears the way for future research about *H. ostrearia* and will help characterise the largely unknown genetics of this species.

The *H. ostrearia* used in this study was isolated in 2018 from an oyster pond in Bouin, France (46°57′12.4″N, 2°02′46.1″W) and deposited in the Nantes Culture Collection (Vona Meleder; Nantes, France) under the name NCC 532. During 21 days, the culture was grown in enriched artificial sea water (*Instant Ocean*, Aquarium systems O ; Harrison et al. 1980 modified by De Brouwer et al. 2002) at 14°C under 300 μm photons/m2/s with a 14h/10h light/dark cycle. On the 20th day of growth, a 1:100 dose of Sigma’s antibiotic antimycotic solution (Sigma-Aldrich, Saint-Quentin Fallavier, France; catalogue#A5955) was added to the culture mix. After 24h, the biomass was collected by filtration (Whatman™ Binder-Free Glass Microfiber Filters, Grade GF/C, pore size of 1.2 μm) and the whole DNA was extracted using the method of Puppo et al. (2017). Extracted DNA was sequenced using the PacBio continuous long reads (PB CLR SEQUEL2) and Illumina MiSeq platforms (TrueSeqv3, 250pe; Genotoul, Toulouse, France), and 85 Gb and 2.36 Gb total read lengths were generated, respectively. Illumina reads were filtered to remove low-quality reads (<Q30), short reads (<75b) and adapter sequences were searched and trimmed by *trimmomatic* 0.39 (Bolger et al. 2014). The *de novo* genome assembly was performed using *Flye* 2.9 (Kolmogorov et al. 2020) and the long PB CLR reads with the following options: −g 100m --meta. Polishing was performed with filtered Illumina reads through three loops of *racon* 1.4.20 (Vaser et al. 2017) and *bwa-mem* v2 2.2.1 (Vasimuddin et al. 2019). The annotation and gene prediction were performed by *Prokka* v1.14.6 (Seeman 2014). Since the mitochondrial genome is circular, *samtools faidx* v1.14 (Danecek et al. 2021) was used to relocate the annotated origin of H-strand replication (OH) as the starting gene (position +1). Genome maps were generated with *Artemis* 18.2.0 (Rutherford et al. 2000).

Phylogenetic trees were created thanks to *NGPhylogeny.fr* pipeline (options trimAl and PhyML+SMS, Lemoine et al. 2019) with the partial *COX1* gene for the mitochondrial and the partial *rbcL* gene chloroplast genomes. The mitochondrial dataset grouped sequences from different diatom taxa: 5 *H. ostrearia* strains, 4 other *Haslea* species, 5 *Navicula* species and 2 external species from the *Eunotia* family. The chloroplast dataset also included different diatom sequences: 3 *H. ostrearia* strains, 5 other *Haslea* species, 8 *Navicula* species and 2 external species from the *Eunotia* family. All sequences were downloaded from NCBI website (https://www.ncbi.nlm.nih.gov/).

The complete circular mitochondrial genome is 38,696 bases long with a GC content (%GC) of 28.66% (36.07% A, 35.26% T, 14.76% C, 13.91% G), close to the *H. nusantara* mitochondrion(36,288 bases, 29.24% GC; Prasetiya et al. 2019). The sequence depth associated with this assembly is 1,325X. The 64 annotated genes were more numerous than *H. nusantara* ones with 3 additional genes, for a total of 39 CDS, 2 rRNA genes and 23 tRNA genes (Figure 1A). The *H. ostrearia COX1* gene, previously partially sequenced (Gastineau et al. 2013), was found complete in this study and annotated as *ctaD* by Prokka. The draft chloroplast genome was 130,200 bases long with 31.04% GC (34.19 A, 34.76% T, 15.31% C, 15.73% G), almost the same as the *H. nusantara* chloroplast (120,448 bases and 31.10%GC ; Prasetiya et al. 2019), with a sequence depth of 2,371X. One hundred and thirty one CDS, 6 rRNA genes, 31 tRNA genes and 1 tmRNA gene were identified within this genome (Figure 1B), quite close to the annotated genes of *H. nusantara* chloroplast. The known *H. ostrearia rbcL* gene, annotated as *cbbL*, and the *psbC* partial gene (Gastineau et al. 2013) were also found and completed in this study. Phylogenetic trees of both mitochondrial and chloroplastic sequences support the hypothesis that *H. ostrearia* strains are very close to each other and have a different evolutionary history from other diatoms or *Haslea* species (Figure 2). However, due to a lack of information about genes in the genus *Haslea*, only the *COX1* and the *rbcL* genes were tested.

**Figure 1:**
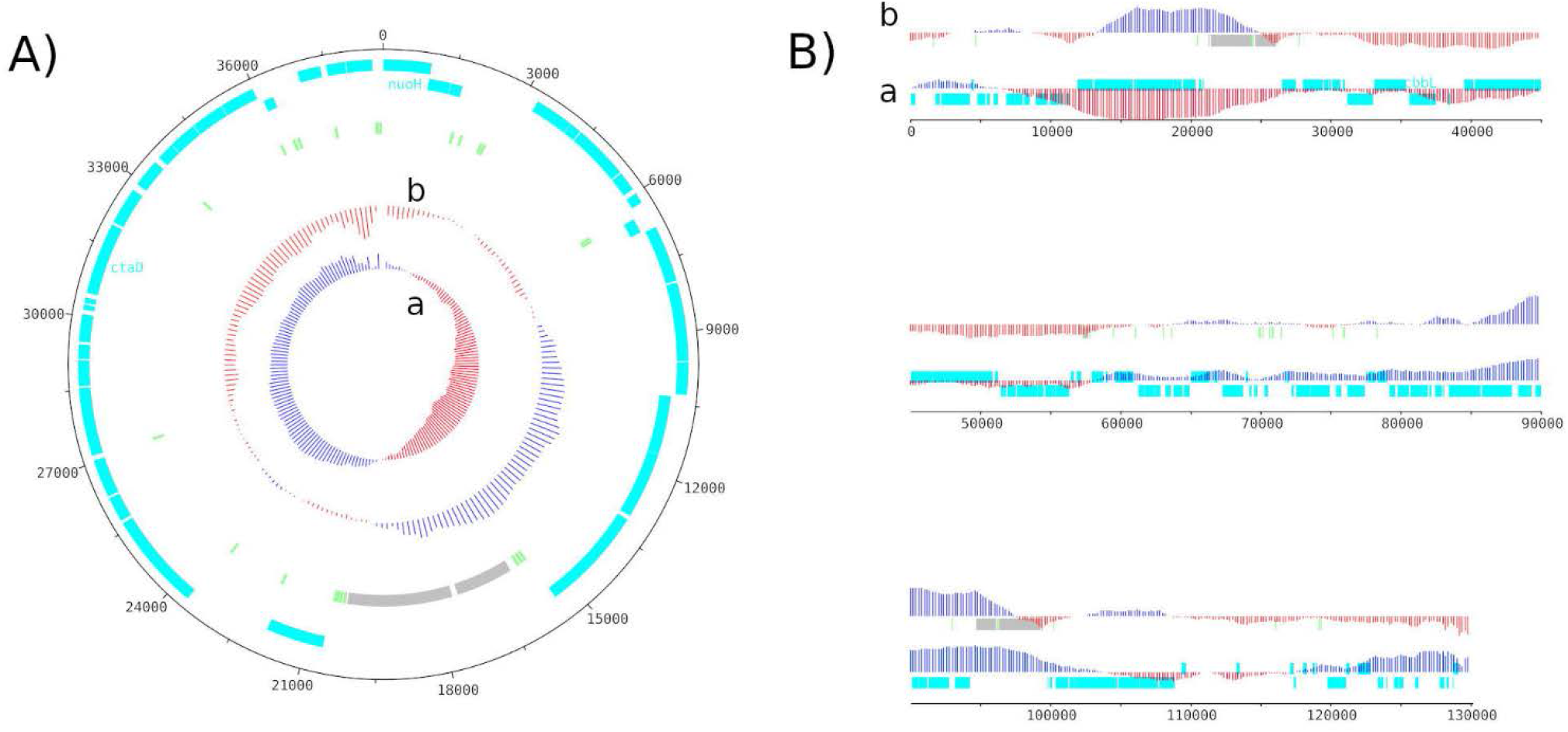
Genome Map of the circular mitochondrial genome (A) and the linear chloroplastic genome (B). The red and blue colors in GC shiew (a) and GC plot (b) show if the value is below or above average, respectively. CDS are shown in light blue, tRNA in light green and the rRNA in light grey.

**Figure 2:**
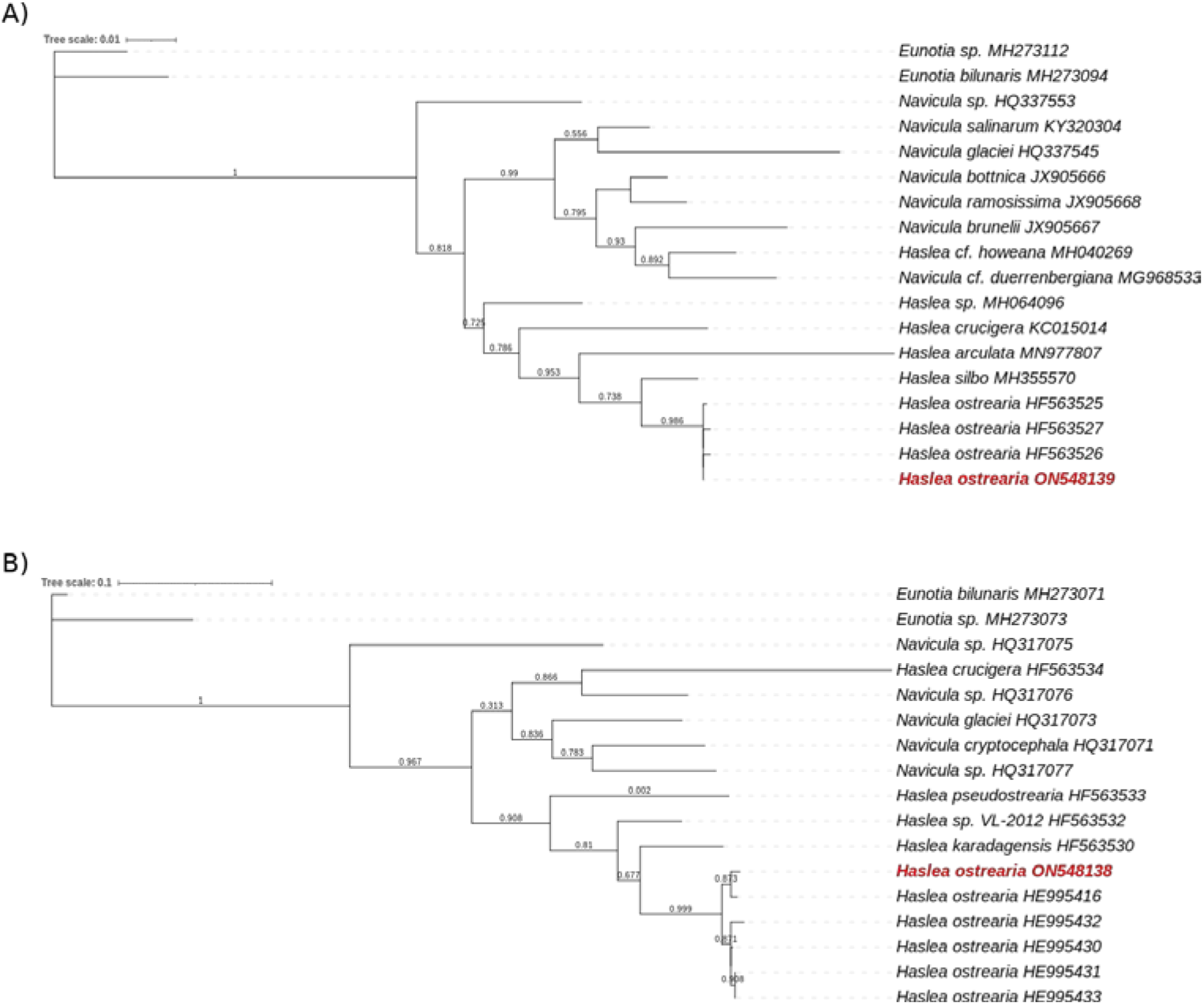
Maximum likelihood phylogenetic trees inferred from COX1 and rbcl genes from, respectevily, *Haslea ostrearia* (in red) and 16 to 18 other diatom chloroplastic (A) and mictochondrial (B) genomes. Numbers near the nodes indicate bootstrap support values.

Even if the chloroplast genome is not completed, it seems that only a few bases are missing since its size and its composition (%GC and genes) are similar to the plastid from the close species *H. nusantara*. To fill the last gaps, further analyses are needed, probably with other sequencing data type. Since all *H. ostrearia* strains for which DNA sequences are available present a close proximity to each other, the mitochondrial and chloroplast genomes may be used as a reference for the species.

## Acknowledgments

We express our sincere thanks to Mr. Christophe Klopp for helping us to reconstruct H. ostrearia genome, and to Mr Bruno Cognie for isolating the strain studied here. All sequencing analyses have been carried out by the Genotoul platform (Toulouse, France).

## Data availability statement

The genome sequence data that support the findings of this study are openly available in GenBank of NCBI at https://www.ncbi.nlm.nih.gov/ under the accession no. ON548138-ON548139. The associated BioProject, SRA and Bio-Sample numbers are PRJNA843895, SRR19450090/SRR19450089, and SAMN28772203 respectively.

